# Lysosomal cathepsin D regulates bone turnover through distinct mode of actions of the autophagy pathways in osteoblasts and osteoclasts

**DOI:** 10.1101/2025.04.09.645406

**Authors:** Fengjie Zhang, Yichen Zhao, Qiling He, Lonnie Schneider, Jianghua Zhang, Chao Wan

## Abstract

Insufficiency in nutrient availability, oxidative stress and autophagy failure are fundamental factors for the decline of bone mass and strength with aging. Accumulating evidence indicates that these factors affect normal autophagosomal or lysosomal activities which are the major force in clearance of aggregated or damaged proteins. Cathepsin D (CtsD), the principal lysosomal aspartate protease and a main endopeptidase, exists in the skeleton during development or homeostasis. However, the molecular and cellular mechanisms of CtsD mediated autophagosome or lysosome function in the skeletal homeostasis remain unclear. In the present study, we showed that deletion of CtsD dramatically decreased bone mass in the 3-week old mutant mice compared with their control littermates as indicated by decreased bone volume (BV), bone volume / total volume (BV/TV), bone surface (BS), trabecular number, trabecular thickness and increase trabecular separation in the microCT analysis. Histomorphometry analysis revealed that the phenotype was characterized by decreased osteoblast numbers, osteoblast surface/bone surface and mineral apposition rate, increased osteoclast numbers, osteoclast surface/bone surface and erosion surface/bone surface. At molecular level, siRNA medicated inactivation of CtsD in MC3T3E1 cells attenuated osteoblastic differentiation and downregulated LC3B expression, which was accompanied by decreased levels of P62, p-Akt and p-GSK3beta in osteoblasts. Intriguingly, inactivation of CtsD in RAW264.7 cells increased osteoclast differentiation with decreased LC3B expression but upregulated P62 level. This was accompanied by alterations in the formation of autophagosome and differential transcription profiles associated with the autophagy pathway during the differentiation of osteoblasts and osteoclasts. The results suggest that CtsD mediated autophagy pathway plays important roles in regulation of bone mass and homeostasis through distinct mode of actions in osteoblasts and osteoclasts, and CtsD may serve as a potential therapeutic target for the maintenance of bone mass.

## INTRODUCTION

Deficiency or mutation of genes encoding lysosomal enzymes or transport proteins causes a group of inherited metabolic disorders - lysosomal storage disorders (LSDs) (1,2). The initial effect of such disorders is accumulation of specific metabolic substrates or precursors inside the endosomal - autophagic - lysosomal system, which subsequently results in cellular dysfunction and impairment of multiple organs (2,3). The most frequently affected organs include brain, viscera, bone and cartilage (4–6). Skeletal pathologies are frequently observed in LSDs, yet the relevance of specific lysosomal enzymes in the regulation of bone turnover remains largely unknown (7,8).

Physiological bone turnover or homeostasis is precisely controlled by the balance of osteoblastic bone formation and osteoclastic bone resorption. Multiple growth factors including transforming growth factor-beta (TGF-β), bone morphogenetic protein (BMP), Wnts, vascular endothelial growth factor (VEGF), hormones such as parathyroid hormone (PTH), estrogen, insulin, insulin like growth factor-1 (IGF-1) (9,10), and transcription factors like Runt-related transcription factor 2 (RUNX2) and hypoxia inducible factor-alpha (HIF-α) have been shown to be involved in the regulation of osteoblast or osteoclast gene programs (11,12). Autophagy, a conserved self-degradative process for balancing sources of energy and in response to nutrient stress, also play important roles in regulation of osteoblast and osteoclast survival, differentiation and function (13–16). Dysregulation of the above regulators impairs the coupling of bone formation and resorption, which consequently leads to metabolic bone disorders such as osteoporosis, osteolysis, and osteopetrosis.

More than 60 hydrolysis enzymes in the lysosomal lumen and above 50 membrane proteins of the lysosome have been identified (17). Among them, cathepsins are the major class of hydrolytic enzymes in the lysosomes. Based on the catalytic mechanisms and structures of the enzymes, the cathepsin family members are categorized into three types: aspartic, serine, and cysteine proteases. Aspartic type cathepsins D and E belong to the A1 peptidase family with human pepsin A as the prototypic enzyme. Serine-type cathepsins A and G belong to the S10 and S1 family, respectively. Cysteine-type cathepsins include cathepsins B, C, F, H, K, L, V, O, S, W, and Z, sharing homology with papain from Carica papaya and are classified as members of the C1 protease family (18). It is indicated that cathepsins B, H, K, L and S are present in osteoclasts, hypertrophic and proliferating chondrocytes at the osteochondral junction of epiphyseal growth plates, and are involved in the extracellular matrix (ECM) degradation during endochondral ossification (19). In bone fracture healing, cathepsins B, H, L and S are mainly related to chondrocyte hypertrophy in cartilaginous callus and bone remodelling (20). Clinical data and genetic mouse models indicate that deficiency of single cathepsin protease causes LSDs, which are often associated with various defects in skeletal development, growth and metabolism, summarized as dysostosis multiplex (21–23). For example, cathepsin K secreted by osteoclasts functions as a potent collagenase to digest collagen in the ECM of the bone. Deficiency of cathepsin K leads to osteopetrosis due to impaired osteoclast resorptive function (21,24,25). Mice with cathepsin L mutation have significantly decreased trabecular, but not cortical, bone volume compared with the wild-type mice (23). These data suggest that cathepsins play important roles in skeletal development and repair.

Cathepsin D (CtsD) (EC 3.4.23.5) is a representative aspartic proteinase in the lysosomes and widely exists in various tissues of mammals. As the principal lysosomal aspartate protease and a main endopeptidase, CtsD functions in the normal activity of autophagysomes and lysosomes (26,27). Early studies identify that mutations in the CtsD gene on chromosome 11p15 cause the most aggressive and earliest onset neuronal ceroid lipofuscinosis (CLN), CLN10 (28,29). The pathological hallmark of all CLN is the intracellular storage of ceroid lipofuscins, primarily in the neurons of the central nervous system (30). The disease-causing mechanisms are further corroborated by studies in mice lacking CtsD (31), sheep with congenital ovine CLN (32), and bulldogs with CtsD mutations (33). CtsD-deficiency mice are born normally but die at postnatal day 26 because of massive intestinal necrosis, thromboembolia, and lymphopenia (34). Tissue phenotype examinations suggest that CtsD is not necessary for embryonal development, but indispensable for postnatal tissue homeostasis (34–36). Interestingly, CtsD deficiency causes extensive accumulation of endogenous alpha-synuclein (α-syn) in neurons, which is associated with impaired macroautophagy and reduced proteasome activity (37). Deletion of CtsD impairs the effective degradation of excess α-syn in dopaminergic cells, resulting in LSD and facilitates α-syn toxicity by its misprocessing (38). In addition, CtsD is also involved in rather non-specific protein degradation in a strongly acidic milieu of lysosomes. CtsD is one of the major proteinases that have the ability to degrade collagens and proteoglycans, the major ECM components of the skeletal tissue (39). Besides the above mentioned, CtsD is also shown to exist in osteoblasts (40), osteoclasts (41,42) and chondrocytes (43) under pathophysiological conditions, yet its functional roles in the homeostasis of the skeleton remain unclear.

In this study, we show that mice lacking CtsD have severe bone loss compared with their control littermates. The phenotype is characterized by decreased osteoblast numbers, impaired osteoblastic bone formation, increased osteoclast numbers and enhanced osteoclastic bone resorption. At molecular level, inactivation of CtsD reveals differential expression pattens of the autophagic pathway gene profiles between osteoblasts and osteoclasts. Inactivation of CtsD in osteoblastic cells downregulates the levels of autophagic markers LC3BII and P62 accompanied by decreased PI3K-Akt-mTOR signals, while inactivation of CtsD in osteoclastic cells shows decreased LC3BII and elevated P62 level accompanied by activation of the PI3K-Akt-mTOR pathway. Our data indicate that CtsD acts as a distinct mediator of the autophagic pathway in regulation of osteoblastic bone formation and osteoclastic bone resorption, which subsequently controls bone homeostasis and bone mass.

## RESULTS

### Inactivation of CtsD decreases bone mass and impairs trabecular microarchitecture

To determine the potential role of CtsD in bone development and remodeling, we generated the transgenic mouse strain with deletion of CtsD. The skeletal phenotypes of the CtsD mutant mice were systemically examined. X-ray analyses of the long bones and lumbar vertebrae showed a decrease in bone mass and density in the *CtsD^−/−^* mice compared with the *CtsD^+/−^* and *CtsD^+/+^* control littermates at 25-day of age (Fig. 1A, B). Consistently, the cortical bones of the femurs and calvariae from the *CtsD^−/−^* mice exhibited the same phenotype (Supplemental Fig. 1A, B). MicroCT measurement on the trabecular bones of the femurs from the *CtsD^−/−^* mutant mice showed a dramatic decrease in bone volume and alterations in trabecular microarchitecture compared with the controls (Fig. 1C). The *CtsD^−/−^* mutant mice had significantly decreased trabecular bone volume, the ratio of bone volume and total volume, bone surface, trabecular number, and thickness, while increased trabecular separation compared with that of the control littermates (Fig. 2D-2I). These data suggest that inactivation of CtsD leads to decreased bone acquisition and the impairment of the trabecular microarchitecture.

**Fig. 1.**
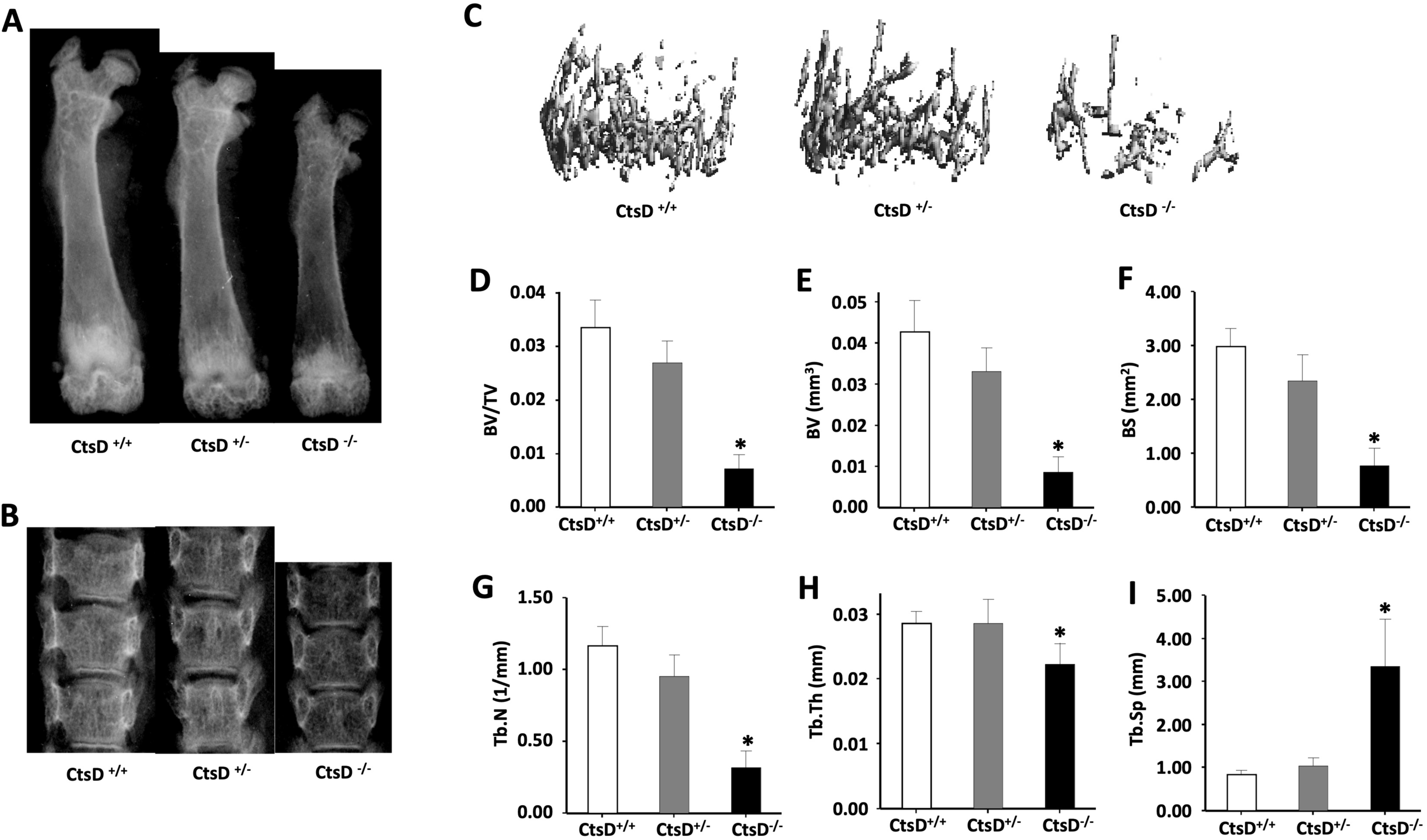
Inactivation of CtsD decreases bone mass and alters microarchitecture of the trabecular bone. The femurs were harvested from the CtsD mutant (*CtsD^−/−^*), heterozygous (*CtsD^+/−^*), and control littermates (*CtsD^+/+^*) at 25-day old, fixed in 10% formalin, and processed for X-Rays and micro-CT analysis. Representative X-ray images of the femurs **(A)** and lumbar vertebrae **(B)** from the *CtsD^−/−^*, *CtsD^+/−^*, and *CtsD^+/+^* mice at 25-day old. The bone density in *CtsD^−/−^* mice is obviously decreased. **(C)** Representative 3D reconstructions of the distal femur. Quantitative analysis shows that mice lacking CtsD have decreased bone volume/total volume (BV/TV) **(D)**, bone volume (BV) **(E)**, bone surface (BS) **(F)**, trabecular number **(G)**, trabecular thickness **(H)**, and increased trabecular separation **(I)**. \**P* < 0.05, n = 6.

**Fig. 2.**
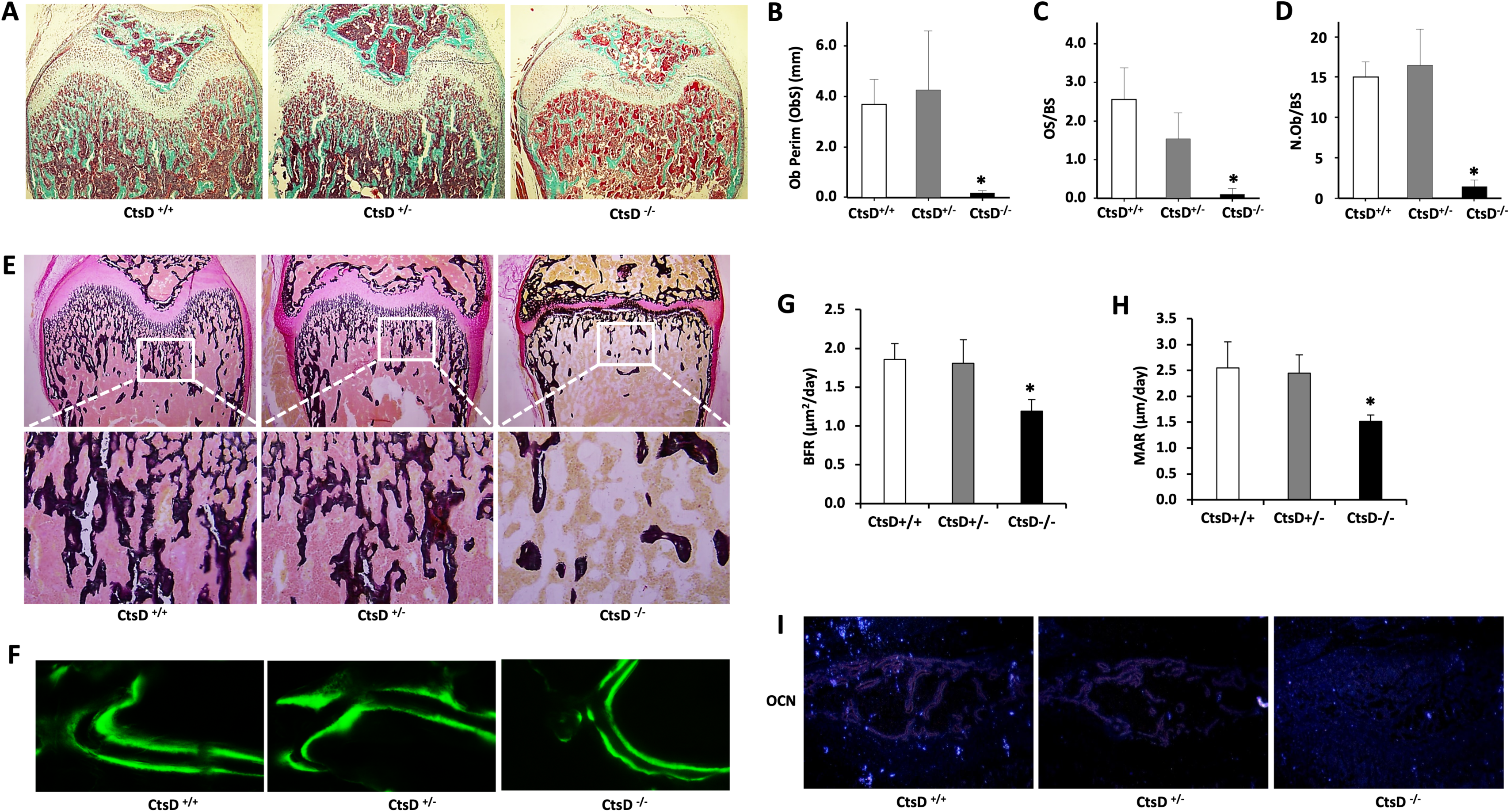
Inactivation of CtsD decreases osteoblast number and impairs bone formation. Non-decalcified or decalcified bone sections were subjected to histochemical staining and bone histomorphometry analysis. **(A)** Representative images of Goldner trichrome staining of the bone sections from the *CtsD^−/−^*, *CtsD^+/−^*, and *CtsD^+/+^* mice. **(B - D)** The bone histomorphometry analysis shows that inactivation of CtsD causes decreased osteoblast perimeter (**B**), osteoblast surface (**C**) and osteoblast number (**D**), indicating decreased osteoblastic function. **(E)** Von Kossa staining reveals dramatically decreased mineralized trabeculae in the *CtsD^−/−^*mutants. **(F)** Representative images of double calcein-labeled sections of the distal femur from the *CtsD^−/−^*, *CtsD^+/−^*, and *CtsD^+/+^* mice are shown. Original magnification, × 400. **(G)** The bone formation rate (BFR) is shown. **(H)** Mineral apposition rate (MAR) calculated in micrometers per day (µm/d) is shown. \**P* < 0.05, n = 6. **(I)** In situ hybridization for osteocalcin (OCN) shows decreased osteoblast differentiation in the *CtsD^−/−^* mutants.

### Inactivation of CtsD decreases osteoblast number and bone formation

We next examined the effect of CtsD on bone formation and mineralization in the *CtsD^−/−^* mutant mice. Inactivation of CtsD decreased osteoblastic bone formation and mineralization as indicated by Goldner trichrome staining (Fig. 2A) and von Kossa staining (Fig. 2B). To further examine the impact of CtsD on bone formation, both the static and dynamic bone histomorphometric parameters were quantified. Mice lacking CtsD showed dramatically decreased osteoblast number (Fig. 2C), osteoblast perimeter (Fig. 2D), and osteoblast surface (Fig. 2E) compared with that of the heterozygous and control littermates. An analysis of double calcein labeling showed a decreased mineral apposition rate (MAR) and bone formation rate (BFR) in the *CtsD^−/−^* mice than that of the heterozygous and control littermates (Fig. 2F-H). In addition, in situ hybridization analysis indicated that the mature osteoblastic marker osteocalcin was dramatically decreased in the *CtsD^−/−^* mutant mice relative to that of the heterozygous and control littermates (Fig. 2I). These data demonstrate that inactivation of CtsD leads to decreased bone formation and mineralization by decreasing both the number and activities of the resident osteoblasts.

### Inactivation of CtsD inhibits osteoblast differentiation and proliferation *in vitro*

In order to exclude the interference from systemic factors of the mutant mice, primary osteoblasts were isolated from calvariae and cultured with the normal growth medium. CtsD knockdown was performed using siRNA approach. An efficient knockdown of CtsD (90%) was obtained following treatment with CtsD siRNA. After incubation with CtsD siRNA and control siRNA for 48 hours, respectively, the cells were further cultured with osteogenic medium. It showed that the cells with inactivation of CtsD had reduced osteoblastic differentiation as indicated by staining for alkaline phosphatase (ALP) (Fig. 3A, C) and ARS staining for mineral deposition (Fig. 3B, D) when the cells were cultured in osteogenic medium for 14 days or 21 days. Real-time PCR analysis showed a significant downregulation in the expression of Runx2, OSX, ALP, and OCN mRNA levels following knockdown of CtsD in osteoblasts compared with the siRNA control group (Fig. 3E-H). In addition, BrdU incorporation assay showed that knockdown of CtsD in osteoblasts decreased cell proliferation (Fig. 3I). These results indicate that inactivation of CtsD inhibits osteoblast differentiation and proliferation *in vitro*, consistent with the *in vivo* phenotypes.

**Fig. 3.**
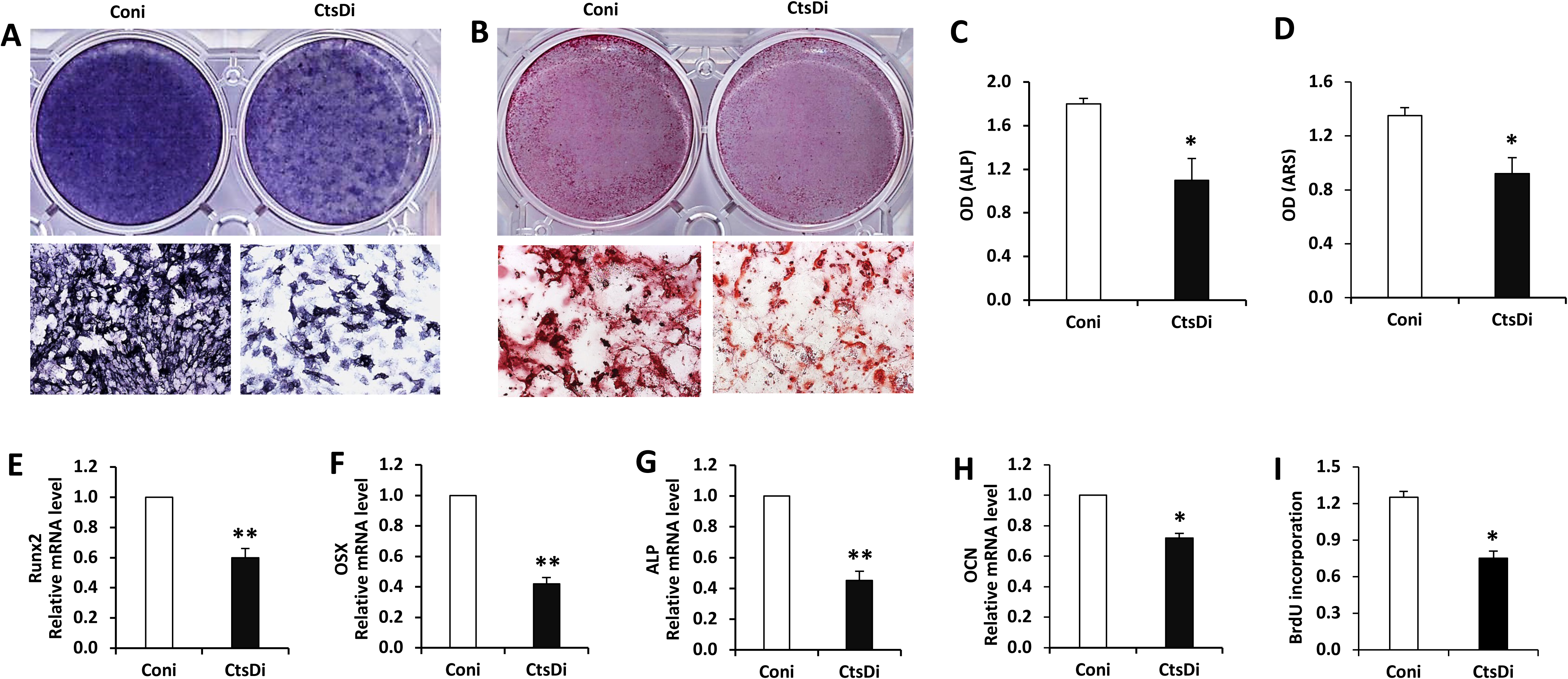
Inactivation of CtsD inhibits osteoblast differentiation *in vitro*. Primary osteoblasts were isolated from calvaria of 3-day old mice and cultured under defined osteogenic conditions. CtsD knockdown was performed using siRNA approach (CtsDi), with control siRNA as controls (Coni). **(A)** Representative images of ALP staining for osteoblasts cultured in osteogenic medium for 14 days. **(B)** Representative images of ARS staining after the osteoblasts cultured in osteogenic medium for 21 days. **(C)** Quantitative integral optical density (IOD) analysis of ALP activity in (A). **(D)** IOD analysis of ARS staining in (B). **(E - H)** Osteoblasts were subjected to siRNA mediated CtsD knockdown, then cultured with osteogenic medium for 7 days. Cells with CtsD knockdown have reduced differentiation potential indicated by reduction in the expression of osteogenic marker genes such as Runx2 (E), OSX (F), ALP (G), and OCN (H). **(I)** BrdU incorporation assay showing decreased proliferation in osteoblasts with siRNA mediated CtsD knockdown. \**P* < 0.05, \*\**P* < 0.01, n = 3.

### Inactivation of CtsD causes lysosomal storage accumulation and reduces p62 levels in osteoblasts

It was indicated that deletion of CtsD caused storage of ceroid lipofuscins in neurons of the brain (28, 29). Here, we also observed lysosomal storage accumulation in osteoblasts of trabecular bone in the *CtsD^−/−^* mutant mice when compared with that of the control littermates as indicated in the TEM analysis (Fig. 4A). We then further examined the potential effect on autophagy in osteoblasts following deletion of CtsD. Confocal immunofluorescence demonstrated that the amount of LC3B positive autophagosomes was dramatically decreased in osteoblastic cells with CtsD knockdown than that of the controls (Fig. 4C), as shown by significantly decreased numbers of LC3B positive puncta (Fig. 4D) and integral optical density (IOD) of LC3B (Fig. 4E). Western blot analysis showed that knockdown of CtsD downregulated LC3B and p62 levels in osteoblasts (Fig. 4F-I).

**Fig. 4.**
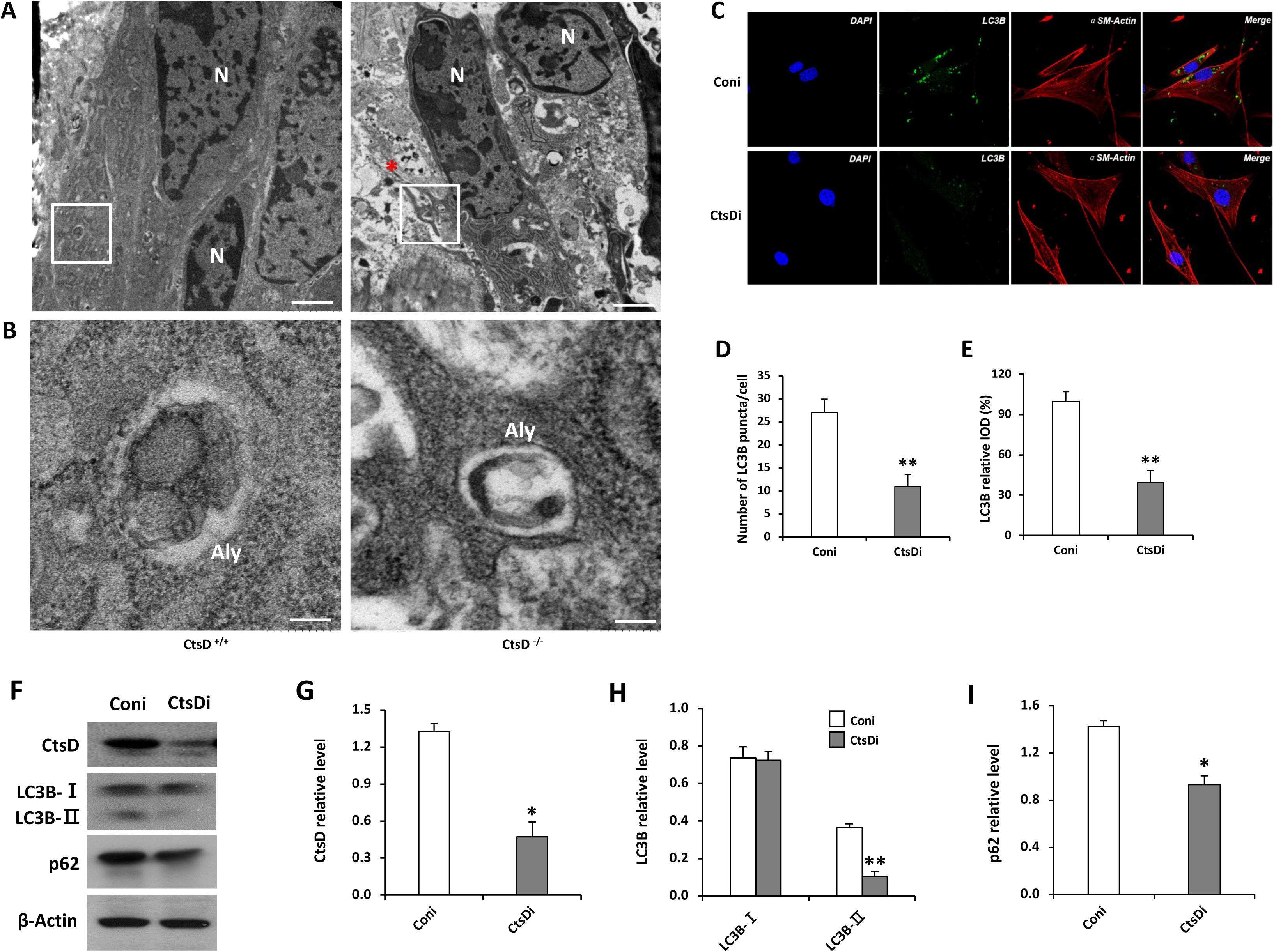
CtsD-deficient osteoblasts displays lysosomal storage accumulation and reduces p62 levels in osteoblasts. **(A)** Representative TEM images showing lysosomal storage accumulation in osteoblasts of the distal femurs from the *CtsD^−/−^* mice at p21 compared with that of the *CtsD^+/+^* mice. The asterisk indicates the lysosomes filled with vacuoles and high electron density materials. Scale bar: 1 μm. **(B)** Higher magnification images of the boxed area in (A) showing altered autolysosome in osteoblasts of the CtsD^−/−^ mice at P21, compared with that of the *CtsD^+/+^*mice. N, nucleus; Aly, autolysosome. Scale bar: 200 nm. **(C)** Confocal immunofluorescence was performed in osteoblasts with siRNA mediated knockdown of CtsD, note that knockdown of CtsD decreased the amount of LC3B positive autophagosomes in osteoblastic cells compared with the siRNA control group. **(D)** Quantitation of the number of LC3B puncta in osteoblastic cells. (**E**) Quantitation of IOD of LC3B in (C). **(F)** Western blot analysis for CtsD, LC3B and p62 in osteoblastic cells following siRNA mediated CtsD knockdown using their specific antibodies. β-actin was used as loading control. **(G - I)** The densitometric quantitation of the immunoblots for CtsD (G), LC3B (H) and p62 (I) in (F). Coni, open bars; CtsDi, solid bars. \**P* < 0.05; \*\**P* < 0.01. n = 3.

To further define the underlying mechanisms, supper array gene profiling for autophagy related genes was performed in MC3T3E1 osteoblastic cells with siRNA mediated knockdown of CtsD. Note that altered autophagy gene profiles were observed in osteoblasts with CtsD inactivation. The most significant changes of 24 genes from the PCR array were selected for further analysis and the hierarchical clustering and expression fold-changes of individual genes were shown in Fig. 5A and 5B respectively. The results indicated that disruption of CtsD led to a remarkable up-regulation of autophagy related 9 homolog B (Atg9b) (4.1-fold) and autophagy-related 16-like 1 (1.25-fold). Interesting, the majority of autophagy related genes were down-regulated in response to CtsD knockdown, particularly, cathepsin S (Ctss) (0.29-fold), phosphoinositide-3-kinase gamma (Pik3cg) (0.3-fold) and microtubule-associated protein 1-3 beta (Map1lc3b) (0.43-fold). Thus, the PCR array data confirmed the effect of CtsD on modulating key genes associated with autophagy in osteoblasts. Further, disruption of CtsD in osteoblasts reduced phospho-PI3K P85, phospho-AKT and phospho-mTOR (Fig. 5C-5I).These results indicate that CtsD plays an important role in regulating osteoblast function during bone growth at least partially mediated by the impairing autophagy and the PI3K/AKT/mTOR signalling process.

**Fig. 5.**
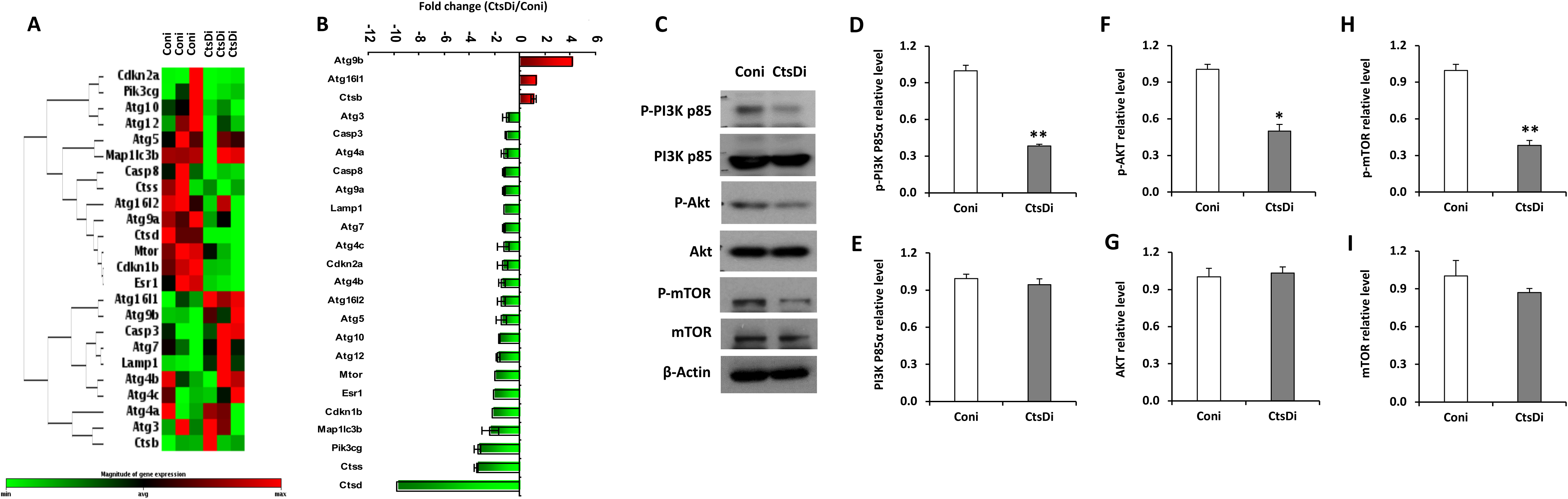
Inactivation of CtsD in osteoblastic cells alters transcription profiles associated with autophagy and inhibits PI3K/AKT/mTOR signalling. **(A, B)** PCR array shows that transcription profiles associated with the autophagy pathway are altered in MC3T3E1 osteoblastic cells with siRNA mediated knockdown of CtsD. Shown is the hierarchical clustering analysis of the PCR array data (**A**) and illustration of the mRNA fold-change of the top three upregulated and 21 downregulated autophagy pathway genes (**B**). **(C)** The knockdown of CtsD in MC3T3 cells was achieved by siRNA approach, and the cell lysates were immunoblotted with antibodies against phospho-PI3K p85, PI3K p85, phosphor-Akt, Akt, phospho-mTOR, and mTOR. Β-actin was used as internal control. **(D - I)** The densitometric quantification of blots for p-PI3K p85, PI3K p85, p-Akt, Akt, p-mTOR, and mTOR in (C) were shown in (D), (E), (F), (G), (H) and (I), respectively. \**P* < 0.05; \*\**P* < 0.01. n = 3.

### Inactivation of CtsD increases osteoclast number and resorptive activity

We next examined the role of CtsD in regulation of osteoclast function. TRAP staining analysis was performed in the femur sections from the *CtsD^−/−^* mutant mice and control littermates at age of 25 days (Fig. 6A). We found that the osteoclast number was dramatically increased in the *CtsD^−/−^* mutant mice compared with that of the *CtsD^+/+^* and *CtsD^+/−^* mice (Fig. 6B). In parallel, the osteoclast surface and erosion surface were also increased in the *CtsD^−/−^* mutant mice (Fig. 6C and 6D). Consistent with the *in vivo* data, an increase in osteoclast differentiation *in vitro* was also observed in osteoclastogenesis of BMMNCs from the *CtsD^−/−^* mutant mice treated with M-CSF and RANKL for 4 days compared with that of the *CtsD^+/+^*control littermates (Fig. 6E). The number of osteoclast (TRAP positive cells with multi-nuclear) and the size of osteoclast were increased in the *CtsD^−/−^*mutant mice than that of the controls (Figure 6F and 6G). Real-time PCR confirmed an elimination of CtsD in osteoclastic cells (RAW 264.7 cells) by siRNA mediated knockdown. The expression of osteoclast marker genes including c-Src, RANK, CA-II, TRAP, MMP9 was dramatically increased than that of the control cells (Fig. 6H - 6L). These data indicate that inactivation of CtsD increases osteoclast differentiation and resorptive activity.

**Fig. 6.**
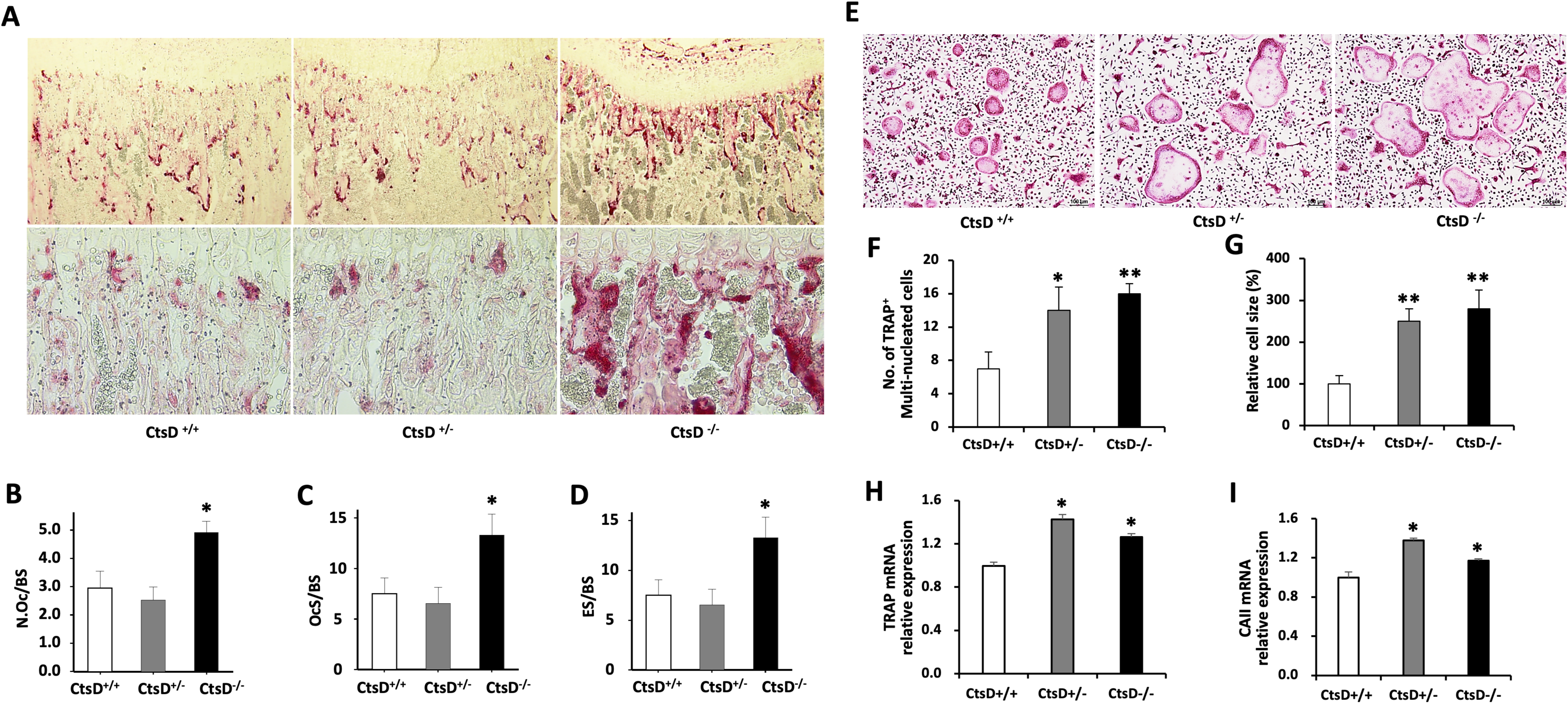
Inactivation of CtsD increases osteoclast number and bone resorption activity. **(A)** TRAP staining shows increased osteoclast numbers and activities in the *CtsD^−/−^*mutants compared with control littermates (*CtsD^+/+^*). **(B - D)** The bone histomorphometry analysis is shown. Inactivation of CtsD causes increased osteoclast numbers (B), osteoclast surface (C) and erosion surface (D). \**P* < 0.05, n = 6. **(E)** BMMNCs were isolated from *CtsD^+/+^* or *CtsD^−/−^* mice and cultured with M-CSF (25 ng/ml) and RANKL (100 ng/ml) for 4 days. TRAP staining was performed to identify multi-nucleated osteoclastic cells. **(F - G)** Quantification of osteoclast cell number (F) and relative cell size in (G) following inactivation of CtsD compared with the control. **(H - L)** Osteoclastic marker genes such as c-Src (H), RANK (I), CAII (J), TRAP (K) and MMP9 (L) mRNA expression are increased in the *CtsD^−/−^* mice than that of the *CtsD^+/+^* mice. \**P* < 0.05, \*\**P* < 0.01, n = 3.

### Inactivation of CtsD causes lysosomal storage accumulation in osteoclasts accompanied by impaired autophagy

To gain further understanding of the molecular mechanisms of the enhanced osteoclastogenesis following deletion of CtsD, TEM analysis of the lysosome, protein analysis and transcriptomic profiling related with autophagy were performed. TEM analysis showed that inactivation of CtsD caused lysosomal storage accumulation in osteoclasts in the *CtsD^−/−^* mutant mice compared with that of the controls (Fig. 7A and 7B), similar to that of the osteoblast. In contrast to osteoblast however, CtsD knockdown in RAW 264.7 cells decreased LC3BII level but upregulates P62 level (Fig. 7C - 7F). Compared with the *CtsD^+/+^* controls, *CtsD^−/−^* mutant mice showed an increase in the number of p62-positive osteoclasts in the trabecular bone of the metaphysis (Fig. 7G - 7H) and upregulated levels of p62 in osteoclasts (Fig. 7G and 7I). To unravel the mechanisms of how disruption of CtsD directly affects autophagy signalling in mouse osteoclast precursor cell line RAW264.7 cells, transcriptomic analysis was performed to compare the transcription level of 84 autophagy related genes, including autophagy machinery components and their associated coregulators, interactors and responsive genes before and after siRNA mediated CtsD knockdown. We found an apparent upregulation of Tgm2 (transglutaminase 2, 1.7-fold), Arkt1 (thymoma viral proto-oncogene 1, 1.6-fold) and Wipi1 (WD repeat domain phosphoinositide interacting 1, 1.5-fold) following CtsD knockdown than that of the siRNA controls. In contrast, Cxcr4 (chemokine CXC motif receptor 4, 3.2-fold), Igf1 (insulin like growth factor 1, 2.5-fold) and Esr1 (estrogen receptor 1, 2.1-fold) were downregulated in response to CtsD knockdown (Fig. 8A and 8B). The results confirmed that knockdown of CtsD altered the transcription profiles of the autophagy pathway in the osteoclastic cells. Further, osteoclastic cells lacking CtsD had higher expression of PI3K p85, AKT, mTOR, and exhibited a significant increase of phospho-PI3K P85, phospho-AKT and phospho-mTOR (Fig. 8C-8I). Taken together, these results reveal that CtsD plays an important role in regulating osteoclasts differentiation and resorptive activity at least partially mediated by the autophagy pathway in osteoclasts.

**Fig. 7.**
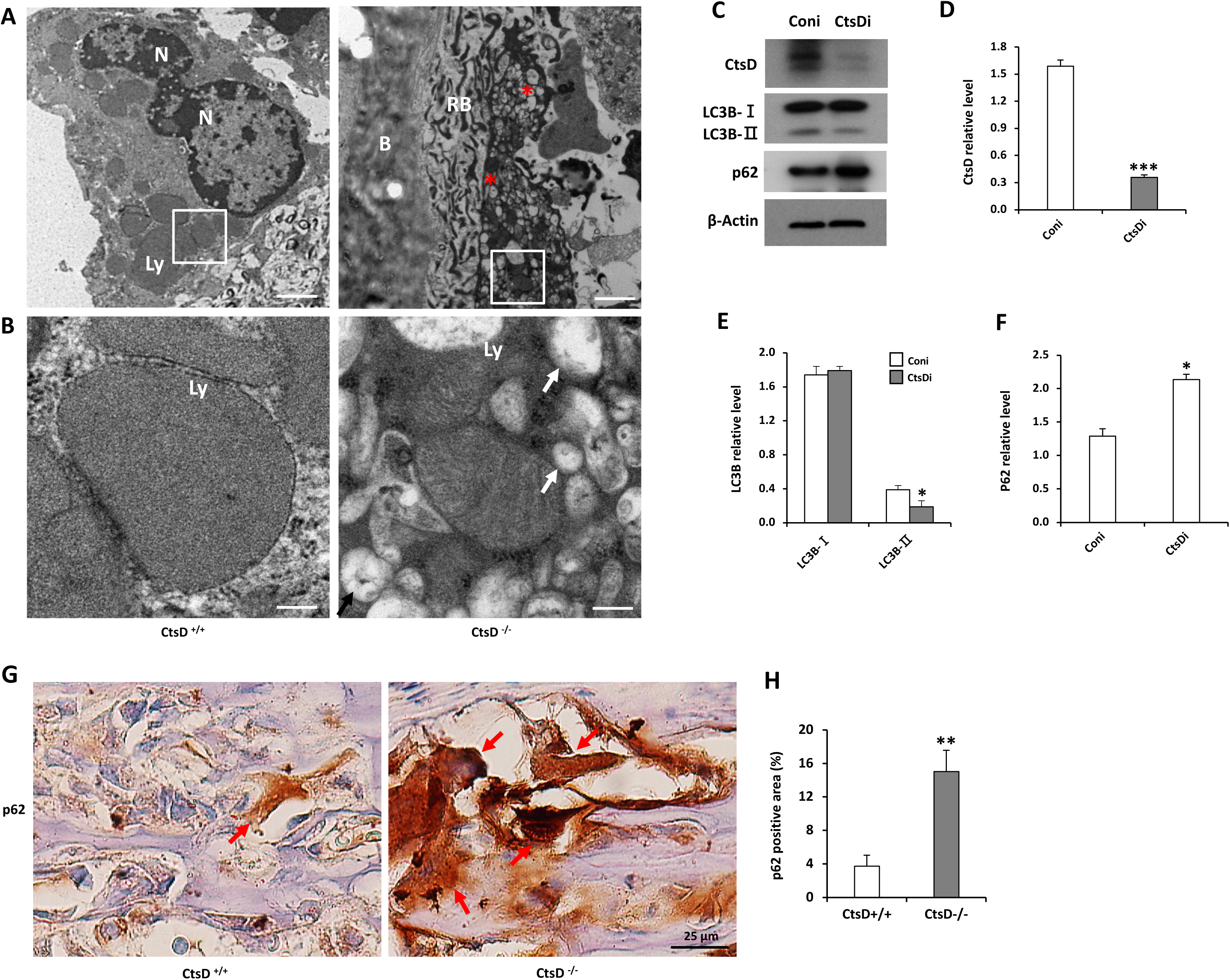
CtsD-deficient osteoclasts exhibit lysosomal storage accumulation and increased p62 levels. **(A)** The representative TEM images showing lysosomal storage accumulation in osteoclasts of the femoral epiphysis from the *CtsD^−/−^*mice, compared with that of the *CtsD^+/+^* mice. The asterisks indicate the lysosomes filled with vacuoles and high electron density materials. N, Nucleus; Ly, Lysosome. B, Bone; RB, Ruffled border. Scale bar: 1 μm. **(B)** Higher magnification images of the boxed area in (A) showing altered autolysosome in the epiphyseal osteoclasts of the CtsD^−/−^ mice at P21, compared with that of the *CtsD^+/+^* mice. The arrows indicate the lysosomes filled with vacuoles and high electron density materials. Scale bar: 200 nm. **(C)** The knockdown of CtsD in RAW264.7 cells was achieved by siRNA approach. Whole-cell lysates were immunoblotted with antibodies against LC3B and p62. Note that CtsD knockdown decreases LC3B-II level and upregulates P62 level. β-actin as internal control. **(D - F)** The densitometric quantification of the immunoblots for CtsD, LC3B and p62 in (C) were shown in (D), (E) and (F), respectively, measuring the ratios of CtsD, LC3B and p62 to corresponding β-actin in the control cells (Coni, open bars) and CtsD cells (CtsDi, solid bars). \**P* < 0.05; \*\*\**P* < 0.001. n = 3. **(G)** Representative histological sections of the distal femurs from 25-day old CtsD+/+ and CtsD−/− mice after immunostaining for p62. Sections were counterstained with hematoxyline. Arrows indicate representative p62 positive osteoclasts. **(H)** The quantification of p62 positive area in (G). \**P* < 0.05; \*\**P* < 0.01. n = 3.

**Fig. 8.**
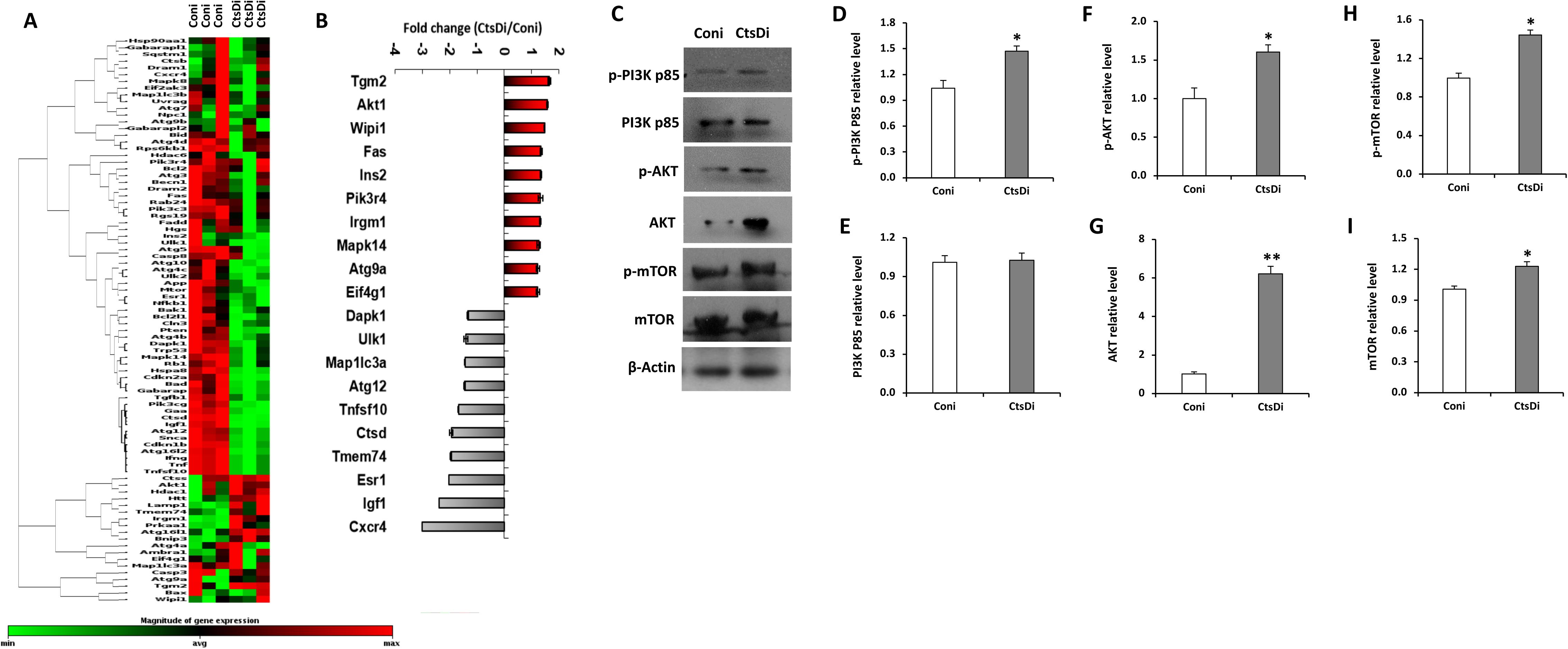
Inactivation of CtsD in osteoclastic cells alters transcription profiles associated with autophagy and activates PI3K/AKT/mTOR signalling. **(A, B)** PCR array gene profiling shows that autophagy gene profiles are altered in RAW264.7 osteoclast precursor cells with siRNA mediated knockdown of CtsD. Hierarchical clustering analysis of the PCR array data (**A**) and illustration of the mRNA fold-change of the top 10 upregulated and 10 downregulated autophagy pathway genes (**B**). **(C)** The disruption of CtsD in RAW264.7 cells was performed with siRNA mediated knockdown, the cell lysates were immunoblotted with antibodies against p-PI3K p85, PI3K p85, p-Akt, Akt, p-mTOR, and mTOR. β-actin was used as internal control. **(D - I)** The densitometric quantification of the immunoblots for p-PI3K p85, PI3K p85, p-Akt, Akt, p-mTOR, and mTOR in (C). \**P* < 0.05; \*\**P* < 0.01. n = 3.

## DISCUSSION

In the present study, we employed a genetic approach to determine the cellular and molecular mechanisms of loss of *CtsD* function in regulation of bone homeostasis. We found that mice with deletion of *CtsD* had significantly decreased bone mass characterized by decreased osteoblast number, osteoid surface, and increased osteoclast number and erosion surface, indicating an impairment of the osteoblastic bone formation and excess osteoclastic bone resorption, leading to imbalance of bone formation and resorption. Mechanistically, the impaired osteoblastic bone formation was caused by the impaired autophagy in osteoblasts, which was associated with suppression of the PI3K-AKT-mTOR signals, altered autophagic gene profiles and decreased p62 level. While the excess osteoclastic bone resorption was related to the altered autophagy in osteoclasts, which was associated with activation of the PI3K-AKT-mTOR signals, alterations of distinctive autophagic gene profiles, and elevated p62 expression. These results suggest that lysosomal CtsD functions is an essential mediator controlling bone turnover or homeostasis through distinct mode of actions in the autophagy pathways in osteoblasts and osteoclasts, respectively.

Histomorphometry analysis in the bone sections from the *CtsD^−/−^* mice showed that the number of osteoblasts and osteoid surface were significantly decreased compared with that of the *CtsD^+/+^* and *CtsD^+/−^* mice, this was consistent with the *in vitro* data that inactivation of CtsD in osteoblasts decreased osteoblast proliferation indexed by BrdU incorporation and osteogenic differentiation indicated by decreased ALP activity and mineral deposition in the ECM of the cultured osteoblasts. This suggests that under physiological conditions, CtsD possesses mitogenic effect on osteoblastic cells and functions as an activator for osteoblast differentiation and mineralization. It is interesting to note that osteoblasts from the *Hyp* mouse, a transgenic strain with loss-of-function mutations of *PHEX* (phosphate-regulating gene with homology to endopeptidases on the X chromosome), produces greater amount of CtsD in the culture media than that of the controls, which suppresses osteoblast mineralization, indicating that overexpression of CtsD might have suppressive effect on osteoblast differentiation (31). These findings suggest that the maintenance of physiological level of CtsD is important for osteoblasts to fulfil their normal function during bone formation.

Another striking phenotype we observed was that mice lacking CtsD had dramatically increased osteoclast numbers and bone resorption than that of the wild-type control and heterozygous mutants. Bone marrow mononuclear cells isolated from the *CtsD^−/−^* mice exhibited enhanced ability of osteoclast differentiation *in vitro* following stimulation with G-CSF and RANKL, suggesting that the enhanced osteoclast differentiation is mainly through the cell-autonomous mechanism. Intriguingly, osteoclasts in the trabecular bone of the *CtsD^−/−^* mice exhibited increased levels of TRAP than that of the heterozygous mutants and wild-type controls. This coincides with the upregulation of the mRNA levels of mature osteoclast markers including TRAP, c-Src, MMP9 and CAII in the trabecular bone tissue harvested from the *CtsD^−/−^* mice. As CtsD is considered as a potent enzyme for digestion of the ECM components (e.g., collagens and proteoglycans) of the skeletal tissue, it was postulated that inactivation of CtsD might reduce the breakdown of the ECM components in the bones, thus leads to increased bone mass. Surprisingly, different from the *CtsK^−/−^* mice, in which osteopetrosis was developed due to inactivation of CtsK, *CtsD^−/−^* mice exhibited severe osteoporosis phenotype. This suggests that CtsD exerts unique functions in regulating osteoclast differentiation and its resorption capacity. However, we could not exclude the alterations of other cathepsin members caused by CtsD inactivation that might also involve in the regulation of bone mass. It is reported that CtsD in pancreatic acinar and inflammatory cells could activate CtsB in a model of experimental pancreatitis (35). To what extend CtsD regulates the other cathepsin members and the underlying mechanisms need to be further explored.

In addition to the reduced proteasome activity, CtsD deficiency impairs macroautophagy and leads to lysosomal dysfunction, which is implicated in the neurodegenerative diseases (36,37). To further define the molecular mechanisms on how CtsD modulates bone formation and resorption through the autophagy pathway, transcriptomic analyses were performed in osteoblastic and osteoclastic cells following siRNA mediated CtsD knockdown. The transcription expression profiles between osteoblastic and osteoclastic cells appeared distinct and unique. For instance, there was a remarkable increase in *Atg9a* expression in osteoblasts but downregulated in osteoclasts. Moreover, both *Atg16l1* and *CtsB* were upregulated in osteoblastic cells but they did not have apparent change in osteoclastic cells. Thus distinct mode of actions of CtsD in the gene profiles related with the autophagy pathway exists, indicating that the CtsD mediated autophagosome/lysosome pathways were differentially regulated in osteoblasts and osteoclasts.

Our data also showed the differential regulation of p62 levels in osteoblastic and osteoclastic cells following the disruption of CtsD. P62, also known as sequestosome-1 (SQSTM-1 or A170), plays important roles in the regulation of cell growth, survival, proliferation and differentiation. A recent study showed that the activity of osteoblasts from the p62−/− mice was reduced compared with those from the wild-type controls, suggesting a role of p62 in regulating the fate of osteoblast precursors toward mature osteoblasts [44]. The role of P62 was complicated in the regulation of osteoclastogenesis. Some previous study demonstrated that osteoclastogenesis was greatly reduced in the bone marrow-derived macrophages from the p62 null mice due to impaired RANK-TRAF6-NFκB signaling [45, 46]. However, a recent study showed that p62 deficiency accelerated osteoclast differentiation [47]. Interestingly, we found that, following knockdown of CtsD, the PI3K/AKT/mTOR signaling was impaired in MC3T3 cells, but dramatically activated in RAW264.7 cells, which coincided with the changes of p62. This suggests that CtsD mediates distinct actions in autophagy in association with its differential regulation of the PI3K/AKT/mTOR signaling in osteoblast and osteoclast.

Taken together, the alterations and imbalance of bone formation and resorption in the *CtsD^−/−^* mice suggest that lysosomal CtsD functions as a key enzyme involved in the regulation of bone homeostasis and bone mass. CtsD may be considered as a potential target for interventions of metabolic bone diseases.

## MATERIAL AND METHODS

### Animals

Homozygous CD knockout mice were obtained through heterozygous breeding. The wild type control, heterozygous and mutant littermates were used for the experiments. The wild type littermates were used as controls. Genotyping was performed on genomic DNA obtained from tail biopsies. All animals were maintained in the Laboratory Animal Services Centre (LASEC), the Chinese University of Hong Kong (CUHK). All the mice were housed in the controlled conditions such as a room temperature of about 22 ± 1°C and a light-dark cycle of 12:12 h. All animal experiments were approved by the University Animal Experimentation Ethics Committee (AEEC), CUHK, and Animal (Control of Experiments) Ordinance from Department of Health, Hong Kong SAR.

### Culture of osteoblasts, BMMNCs and cell lines

Osteoblasts were isolated from long bones of 3-4 weeks old mice by serial digestion in 1.8 mg/ml of collagenase type I solution (Worthington Biochemical Corp.). BM was flushed out and the bone was cut into small pieces and then digest in 10 ml of digestion solution for 20 min at 37°C under constant agitation. The digestion solution was then collected, and the digestion was repeated for additional 4 times. Digestion solutions containing the osteoblasts were pooled together. After centrifugation, osteoblasts were obtained and cultured in αMEM containing 10% FBS and 1% penicillin/streptomycin at 37°C in a humidified incubator supplied with 5% CO_2_. Bone marrow mononuclear cells (BMMNCs) were obtained from *CtsD^+/+^*or *CtsD^−/−^* mice, and culture under defined conditions for induction of osteoclast differentiation with RANKL (100 ng/ml) and M-CSF (25 ng/ml). siRNA mediated knockdown of CtsD was also achieved in MC3T3-E1 pre-osteoblastic cell line or RAW264.7 pre-osteoclast cell line.

### Histological and histomorphometry analysis

Histological examination of the long bones was performed at defined time points of the mice following birth. Specimens were stripped off soft tissues, fixed in 4% paraformaldehyde and decalcified with 10% EDTA. Specimens were embedded in paraffin, sectioned and mounted onto slides. Trichrome staining and H&E staining was performed for histological examination using standard methods. In a parallel study, femur or tibia bone samples was processed to plastic embedding for non-decalcified bone sections. Quantitative histomorphometry analysis was performed using the OsteoMeasure system (OsteoMetrics). Static parameters including osteoblast number, osteoblast surface, osteoid surface, osteoclast number, osteoclast surface (or erosion surface) were calculated, and dynamic parameter mineral apposition rate (MAR) was examined by injection of two sequential doses of calcein (8 mg/10 ml sterile saline) delivered in a total of 0.25 ml 3 and 10 days before death. Mineralization of the trabecular bone was visualized by von Kossa staining on the non-decalcified bone sections.

### TRAP staining

Tartrate-resistant acid phosphatase (TRAP) staining was performed to examine osteoclast activity or function by using the commercially available kit (387A-1KT, Sigma-Aldrich) according to the manual protocol.

### Micro-CT analysis

The bones were dissected free of soft tissues, fixed in 10% neutral buffered formalin for 48 h, and analyzed by a high-resolution mCT imaging system (MicroCT40; Scanco Medical). The scanner was set at a voltage of 70 kV and a current of 113 µA. Direct calculations of histomorphometric parameters were performed with 3D reconstruction software, including bone volume /total volume (BV/TV), trabecular thickness (Tb.Th), trabecular separation (Tb.Sp), and trabecular number (Tb.N).

### Quantitative real time-PCR

Total RNA was extracted from osteoblasts using the RNeasy Mini Kit (Qiagen). First strand cDNA was synthesized from 1 μg of total RNA using Primescript RT master mix (Takara). Quantitative real-time PCR was performed with SYBR Premix Ex Taq (Takara) in ABI Fast Real-time PCR 7900HT System (Applied Biosystems). All samples were performed in triplicates. β-actin was amplified in parallel as an endogenous control. Data were averaged and normalized to endogenous β-actin reference transcripts.

### Transcriptomic profiling

Autophagy related transcriptomic profiling was performed using mouse autophagy RT^2^ Profiler PCR Array to examine the expression of 84 genes (Qiagen). siRNA mediated CtsD gene knockdown in MC3T3 cells and RAW264.7 cells were performed following the established protocol. Total RNA was extracted, reverse transcribed and PCR amplified according to manufacturer’s instructions. Data were analyzed using online SABiosciences RT2 Profiler PCR Data Analysis software available at manufacturer’s webpage.

### Western blot

Cells were washed three times with ice-old PBS, whole cell lysate was obtained by cell lysis buffer in the presence of a protease inhibitor cocktail. Protein concentration was examined by BCA protein assay (Pierce). 50 µg of total protein was denatured, resolved in 10% or 12.5% SDS-PAGE mini-gel, and electrophoretically transferred onto an Immobilon-P membrane (PVDF, Millipore). The membrane was blocked with 5% non-fat dry milk and hybridized with antibodies against CtsD (Santa Cruz, CA, USA), LC3B, P62, phospho-Akt, Akt, phosphor-PI3K p85, PI3K p85, phosphor-mTOR, mTOR and β – actin (Cell Signaling, USA) at 4°C overnight. Membranes were washed extensively with 0.1% Tween-20 in TBS, and then incubated with horse-radish peroxidase conjugated secondary antibody (Molecular Probes) in 1:10000 dilution at room temperature for 1 h. Protein expression signals were detected with the Super Signal West Femto Maximum Sensitivity Substrate (Thermo Scientific, USA) and developed on a x-ray film (Roche). β - Actin was used as internal control.

### Immunohistochemistry and immunofluorescence staining

Whole femurs or tibiae were isolated from mice, fixed with 10% formalin at 4°C for 5 days and decalcified in 10% EDTA at pH 7.4 for 3 weeks. Thereafter tissues were embedded in paraffin and sections were cut for standard hematoxylin-and-eosin stain (Sigma). Immunohistochemical staining was performed following standard protocol. Sections were incubated with specific primary antibodies including anti-P62 (1:100, Cell Signaling, USA), and color development with diaminobenzidine was performed by using Histostain Plus Kit according to the manufacturer’s instructions (Thermo Scientific, MA, USA). The quantification of image was performed with Image J software (NIH, USA). For immunofluorescence staining, the cells were washed with PBS, then fixed with 4% PFA, permeabilized by 0.5% Triton X-100 for 5 min, and pre-blocked with 3% BSA for 1 h. Immunofluorescence staining was performed by an incubation with anti-LC3B (1:100, Cell Signalling, USA) or SMA antibody (Thermo Scientific, MA, USA) at 4°C overnight, followed by secondary antibodies Alexa-488 anti-rabbit or Alexa-647 anti-mouse (Thermo Scientific, MA, USA) for 1 h at room temperature. Cells were washed extensively with 0.1% tween-20 in PBS and mounted with DAPI nuclear stain (Vector Lab). Fluorescent signal was visualized by Olympus FV1200 confocal microscope (Olympus). The image was processed by FV10-ASW 1.7 software (Olympus).

### Alkaline phosphatase staining and Alizarin red S staining

Cells were washed by PBS and then fixed with 10% formalin for 15 min. Thereafter, cells were stained with 1-Step NBT/BCIP solution (Sigma-Aldrich) at 37°C for 10-30 min or until desired colour was developed. Staining solutions were removed after color development and washed extensively with distilled water. The images were scanned or taken a photo in microscope. For Alizarin red S staining, was performed by adding 500 µl of Alizarin red S solution to the wells. After incubation at 37°C for 15 min, the wells were washed three times with distilled water. The image were scanned, and then observed by phase-contrast microscopy. The stained areas were quantified using Image J software.

### Transmission electron microscopy (TEM)

The epiphysis and metaphysis bones were prepared for electron microscopy analysis as established procedure. The bones were shaken in a primary fixative solution containing 2% glutaraldehyde, 0.05 M sodium cacodylate buffer (pH 7.4) with 0.7% ruthenium hexammine trichloride (RHT) (Polysciences) for 3 hours at room temperature, then washed three times for 10 minutes with 0.1 M sodium cacodylate buffer (pH 7.4). Post fixation was carried out with 1% osmium tetroxide in 0.1 M sodium cacodylate buffer (pH 7.4) with 0.7% RHT for 2 hours at room temperature. The samples were washed three times for 5 min with the same buffer. Following fixation, samples were dehydrated in ethanol and washed with propylene oxide twice for 15 minutes at room temperature. The tissue was then subjected to propylene oxide with increasing concentration of epon (30%, 50% and 75%, each overnight at room temperature) and then 100% epon for 5 hours before polymerization. Ultrathin sections (80 nm) were examined with a transmission electron microscope.

### Statistics

The quantitative real-time PCR, ELISA experiments and other quantitative analyses were repeated at least three times unless otherwise stated. Comparisons were made by using Student’s *t-*test or one-way ANOVA according to the experimental design. Results were expressed as mean ± standard deviation (SD). *P* < 0.05 was considered as statistical significance.

## Acknowledgments

We are grateful for the assistance of the Core laboratories at School of Biomedical Sciences, the Chinese University of Hong Kong, and Histomorphometry and Molecular Analysis Core at the University of Alabama at Birmingham. We appreciate Dr. Paul Saftig at Christian Albrechts University of Kiel, Germany for the support on using the transgenic mouse strain. We appreciate the technical support from Dr. Wing Pui Tsang at School of Biomedical Sciences, The Chinese University of Hong Kong. This work was supported by the Hong Kong Research Grant Council General Research Fund (14115414), the InnoHK initiative of the Innovation and Technology Commission of the Hong Kong Special Administrative Region Government, Shenzhen Virtue University Park Laboratory Support Special Fund (YFJGJS 1.0) and the Basic Research Fund (2021Szvup150), Shenzhen, PR China.

